# Mitochondrial genomes of Pleistocene megafauna retrieved from recent sediment layers of two Siberian lakes

**DOI:** 10.1101/2023.06.16.545324

**Authors:** PA Seeber, L Batke, Y Dvornikov, A Schmidt, Y Wang, KR Stoof-Leichsenring, KL Moon, SH Vohr, B Shapiro, LS Epp

## Abstract

Ancient environmental DNA (aeDNA) from lake sediments has yielded remarkable insights for the reconstruction of past ecosystems, including suggestions of late survival of extinct species. However, translocation and lateral inflow of DNA in sediments can potentially distort the stratigraphic signal of the DNA. Using three different approaches on two short lake sediment cores of the Yamal peninsula, West Siberia, with ages spanning only the past hundreds of years, we detect DNA and identified mitochondrial genomes of multiple mammoth and woolly rhinoceros individuals—both species that have been extinct for thousands of years on the mainland. The occurrence of clearly identifiable aeDNA of extinct Pleistocene megafauna (e.g., > 400K reads in one core) throughout these two short subsurface cores, along with specificities of sedimentology and dating, confirm that processes acting on regional scales, such as extensive permafrost thawing, can influence the aeDNA record and should be accounted for in aeDNA paleoecology.

## Introduction

Sedimentary deposits constitute highly valuable archives of past ecosystem changes as they contain dateable layers with organismic remains including ancient DNA (aDNA). Such remains are typically assumed to represent the ecosystem of the time around which the respective stratum was deposited. aDNA from sediments has yielded remarkable insights regarding paleoecology, phylogeography, and extirpation and extinction events of keystone taxa such as mammoths (Haile et al., 2009; Boessenkool et al., 2012; Graham et al., 2016; Murchie, Monteath, et al., 2021). Based on such ancient environmental DNA (aeDNA), a recent study proposed that the woolly mammoth (*Mammuthus primigenius*) may have survived in Eurasia for much longer than previously assumed, as the authors retrieved mammoth DNA sequences in sediment layers that were approximately 4.6–7 thousand years (kyr) younger than the most recent mammoth fossils (Wang et al., 2021); however, in response to this interpretation, Miller and Simpson (2022) opined that these results may be more likely due to taphonomic processes leading to release of aeDNA from the remains of long-dead organisms from permafrost, where it is well preserved.

Conclusions derived from aeDNA isolated from sediment cores rely on the stratified nature of the remains in question and dating of the respective layer by radiometric methods. However, in theory, various physical and geochemical processes such as translocation of DNA through sediment strata (Haile et al., 2007), re-deposition of older sediment carrying DNA of extinct organisms (Arnold et al., 2011), and preservation bias (Boere et al., 2011) can distort the biological signal of aeDNA and thus bias the accuracy of allocation of taxa to specific time periods (Armbrecht et al., 2019). For lake sediments, deposited under aquatic conditions, studies have suggested that leaching is not a concern (Parducci et al., 2017), but it has been observed in soils and cave sediments (Haile et al., 2007). The question of obtaining last appearance dates of extinct taxa using aeDNA in dated sediment layers (Haile et al., 2009) is under discussion, but the surprisingly young records published so far still date to multiple thousands of years before present and thus lie within a timeframe of possible late survival.

## Results and Discussion

In 2019, we retrieved short subsurface sediment cores from two Arctic thermokarst lakes (LK-001 and LK-007, located approximately 5 km apart, over massive permafrost; **Table 1**; Dvornikov et al., 2016) on the Yamal peninsula, Siberia, to extract DNA and assess changes in mammal abundances in the Arctic over the past decades and centuries. From lake LK-001, we collected a secondary core which was sliced in the field at 1-cm steps for Pb^210^ radiometric dating, which indicated that the sediments at the top of this core were deposited recently, and that the core spanned the past few centuries (**Table S7**). The cores for DNA extraction were closed in the field immediately after retrieval and were then transported to the dedicated aDNA laboratories of the University of Konstanz, Germany. In this lab facility, established in 2020, no other samples from the Arctic or from any large mammals had been processed previously. From core LK-001, we isolated DNA and produced genomic double-stranded libraries from 23 samples, from 1.5 to 80 cm core depth (Supplement section 1), according to standard procedures. The core was opened and all subsequent steps until index PCR setup were carried out under customary aDNA laboratory conditions. In particular, the core opening facilities and the lab are located in buildings separated from the downstream molecular genetic analyses, the ventilation of the aDNA lab is based on a HEPA filter system and positive air pressure, and the lab is subjected to nightly UV radiation. Work in the lab is conducted under strict aDNA precautions, adhering to established aDNA protocols (Fulton and Shapiro, 2019). We enriched the libraries for mammalian DNA using a custom RNA bait panel produced from complete mitogenome sequences of 17 mammal species that currently or previously occurred in the Arctic (adapted from Murchie, Kuch, et al., 2021). The enriched libraries were sequenced, and we mapped the sequences against a database of 73 mammal mitogenomes, followed by BLASTn alignment against the complete NCBI nt database. We thus retrieved mitogenomic sequences of mammals that were expected during the age range covered by the core (**Table S9**), e.g., reindeer (*Rangifer tarandus*), Arctic lemming (*Dicrostonyx torquatus*), and hare (*Lepus*); however, throughout the entire core, there were abundant sequences of two species that have been extinct for several thousand years, i.e., mammoth (*Mammuthus primigenius*) and woolly rhinoceros (*Coelodonta antiquitatis*). Twenty-two of the 23 LK-001 libraries produced > 1,000 reads, each, assigned to *Mammuthus*, with read counts ranging from 2,852 to 72,919 (mean 21,140 ± 17,296). Negative controls (extraction and library blanks) did not produce any reads assigned to mammals. In the sample with the highest *M. primigenius* read counts (31.5 cm depth, dated to 81 years), the coverage of the reference mitogenome (NCBI accession NC_007596.2) was 95.3%, (434 (± 213)-fold). Across all samples, 465,080 reads assigned to mammoth were produced, with 98.3% coverage (2,762-± 1,176-fold). Read lengths ranged from 28 to 289 bp (mean 100 ± 44 bp; **Fig. 1**). The number of woolly mammoth reads decreased from lower samples towards the top of the core (**Fig. 1**). Signatures of post-mortem DNA decay were comparably minor (**Fig. 1**), with reference to an *M. primigenius* genome downloaded from NCBI (accession NC_007596.2), and mapping suggested that the sequences throughout the core originated from multiple individuals. Further analyses of the three libraries with the most mammoth reads using mixemt (Vohr et al., 2017) identified a number of mitochondrial haplogroups in the sequences from the core, suggesting that they originated from a multiple individuals (**Fig. 2**.). The haplogroups identified were known to occupy the region, and it seems likely the sequences reflect a history of mammoth occupation at the core site. Twelve of the 23 libraries produced > 100 reads, each, assigned to woolly rhinoceros, with a total of 2,737 reads and a cumulative coverage of 44% (assembled to NC_012681.1 2). Further analyses are mostly focused on the mammoth sequences as these occurred in substantially higher numbers.

**Table 1.** Sediment cores retrieved from two lakes on the Yamal peninsula, Siberia.

| Lake | Coordinates | m above sea level | Area | Water depth | Core length |
| --- | --- | --- | --- | --- | --- |
| LK-001 | 70°16'45.6" N, 68°53'02.8" E | 28 | 38 ha | 17 m | 80 cm |
| LK-007 | 70°16'02.8" N, 68°59'35.7" E | 36 | 39 ha | 14 m | 75 cm |

**Fig 1.**
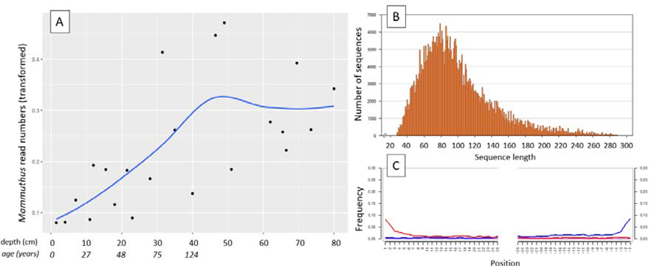
aDNA of *Mammuthus* in recent lake sediments. (**A**) Read counts assigned to *Mammuthus* (square-root-transformed proportion of the respective number of raw reads per library) after hybridization capture enrichment of aeDNA of core LK-001 (shown are results of 22 libraries; one library was excluded as it did not produce any reads assigned to mammals); square-root transformation of percentage. Indicated are sample depths (in cm; 1.5 to 80 cm) and approximate ages as per ^210^ Pb chronology (**Table S7**; to a maximum depth of 39.5 cm). The solid line indicates the general trend. Across the 22 libraries: (**B**) Fragment length distribution and (**C**) damage patterns (red indicates C-to-T transitions, blue G-to-A transitions. the Y-axis indicates the percentage of positions with a nucleotide change, the X-axis indicates the position along the fragment).

**Fig 2.**
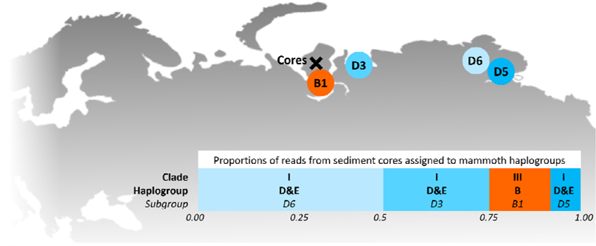
>. Locations of the sediment cores of the present study (Yamal peninsula, Siberia) and previously retrieved mammoth remains and their haplo(sub)groups (**Table S6**). The bar chart indicates a maximum-likelihood estimate of the haplogroup proportions derived from the reads from the three sediment core libraries with the most mammoth reads.

As these results were entirely incongruent with the temporal occurrence of these two species, we employed several methods to confirm their validity, i.e., conventional PCR and Sanger sequencing of a mammoth cytochrome c oxidase subunit I (COI) fragment, mammal metabarcoding, and droplet digital PCR (ddPCR) of a mammoth cytochrome b fragment. We performed these analyses on core LK-001 and, to evaluate whether this is a locally isolated phenomenon, on a second short core, LK-007, from a nearby lake, which had been opened and processed under the same conditions indicated above to prevent contamination. Additionally, we screened the core LK-001 for plant macrofossils, of which three were sent for ^14^C AMS dating.

Conventional PCR and Sanger sequencing confirmed amplification of a COI fragment of *M. primigenius*. Mammal metabarcoding produced mammoth sequences (74 bp) in 13 samples of core LK-001 and 9 samples of core LK-007. In the LK-001 core, the highest *M. primigenius* read count occurred at 31.5 cm (2,992 reads), and in the LK-007 core at 26 cm *M. primigenius* (3,580 reads). ddPCR produced *M. primigenius* sequences in 14 samples of each core. Metabarcoding and ddPCR patterns across the cores were similar, although not completely congruent, as ddPCR appeared to be more sensitive (**Fig. S1**). The dates retrieved by radiocarbon dating were not congruent with the initial age inference suggested by Pb^210^. While the lowest and topmost sample (with ages of 1,547 ± 228 and 339 ± 79 uncal. yrs BP respectively) suggest relatively young ages and agree in their temporal succession, the middle sample, at 51 cm, yielded a radiocarbon age of 8,677 ± 132 yrs.

The mammoth was abundant throughout most of Eurasia during the Pleistocene, but populations declined at the end of the Pleistocene, with the species going extinct in the mid-Holocene (Nogués-Bravo et al., 2008). The youngest fossils from mainland Siberia have been dated to 9,650 years (Stuart et al., 2002). The woolly rhinoceros was a cold-adapted megaherbivore, which was abundant from western Europe to north-east Siberia during the Middle to Late Pleistocene (Kahlke and Lacombat, 2008). This species was predominantly grazing, probably resorting to browsing only due to seasonal vegetation restrictions (Kahlke and Lacombat, 2008; Rey-Iglesia et al., 2021; Stefaniak et al., 2021). The reasons underlying the extinction of this species at ca 14 ka BP are not entirely clear, but it is largely attributed to sudden climate warming and subsequent habitat unsuitability due to vegetation changes (Stuart and Lister, 2012; Lord et al., 2020; Wang et al., 2021), likely coupled with human influence (Fordham et al. 2022), in the Bølling-Allerød interstadial warming.

The present data, which implies frequent and abundant Pleistocene megafaunal DNA throughout a sediment core deposited in the lake over the past centuries suggests that physical processes, rather than presence of live organisms, are responsible for the recovery of this DNA. While not in itself fully conclusive, our data suggests the source of the DNA of Pleistocene mammals from older permafrost deposits, either from a carcass or from sedimentary materials carrying the DNA. The numbers of mammoth sequences increased with depth downcore, with comparably low abundances over the most recent 11 cm of the core (aged < 30 years), pointing to a decrease of input of this DNA through time. The apparently rather limited extent of aDNA damage in the mammoth sequences suggests that the source of this DNA has been preserved exceptionally well, which also suggests an origin from permafrost, and the specific dynamics of thawing and re-deposition of material in the study area offer an explanation.

Here, the active thermokarst likely began during the climatic optimum of the Holocene (9000–3000 BC; Savvichev et al., 2021). The LK-001 and LK-007 lake basins are interbedded mainly into the IV^th^ marine plain, formed in the Kazantsevskaya, Marine Isotopic Stage 5. The permafrost of these lake catchments was formed no earlier than 70—60 kyr BP after the Kazantsevskaya transgression in more sub-aerial conditions and with mean annual temperatures 6 to 7 °C lower than modern temperatures (Baulin et al., 1981). As this area was under coastal-marine conditions for a long period, these lake basins may be paleo-marine remnants, or they were formed later as a result of thermokarst over the segregated or tabular (massive) ground ice during the Holocene climatic optimum (Kachurin, 1961). The area is also subject to abrupt permafrost thaw (thermo-denudation), resulting, for example, in the formation of retrogressive-thaw slumps (thermocirques) and the transport of a large amounts of thawed terrestrial material into the lake water (Dvornikov et al., 2018). Such abrupt permafrost thaw processes normally appear adjacent to lakes and can form specific geomorphological elements,. i.e., thermo-terraces (Kizyakov et al., 2006) within lake catchments and lakes; they are normally polycyclic processes appearing due to the extension of a seasonally thawed layer (active layer) up to the top of massive ice and more favorable thermal conditions within the existing forms (Leibman and Kizyakov, 2007; Kokelj et al., 2009). Traces of past thermo-denudation can be observed within both lake catchments. In catchments of five neighboring lakes, large retrogressive-thaw slumps appeared in recent years (2012-2013) accompanied by the thaw and lateral transport of modern and Late Pleistocene deposits into lakes. Additionally or alternatively, thermo-erosion of upper geomorphological levels and transport via stream networks could transport ancient material into the modern lacustrine sediments. However, the two studied lakes are headwater lakes (with outlet, no apparent inlet) and this option can only be considered in terms of small thermo-erosional valleys within the catchments.

An alternative mechanism for the redistribution of Late Pleistocene material in the sediments is related to subcap methane emission (bubbling) from degrading permafrost beneath the lake bottom. In-lake bubbling can be observed in a circum-Arctic scale: in North-East Siberia, Alaska and Canada. This is common especially in lakes with a depth exceeding two meters, which do not freeze entirely up to the bottom in winter, leading to the formation of a talik (a layer of year-round unfrozen ground that lies in permafrost area). The expansion of the talik may further trigger subcap methane emission, which can reach 40-70 kg yr^-1^ of pure (94%–100%) methane in neighboring lakes (Kazantsev et al., 2020). The constant methane seepage doesn’t allow ice to be formed in winter (whereas the normal winter ice thickness is approximately 1.5 m) and can potentially disturb the stratigraphy of lake sediments. Additionally, dramatic emissions of methane can form craters in terrestrial and lacustrine environments (Dvornikov et al., 2019). In this case Late Pleistocene sediments will well be re-distributed within the water-body and the entire stratigraphy will be mixed. In the case reported here, the simultaneous finding of a Pb^210^ chronology indicating recent sediment deposition and of plant macrofossils that dated to > 8000 yrs BP in a sample from 36.5 cm, suggest lateral input of ancient material, including the mammalian DNA, putatively related to permafrost thawing processes. Numerous studies on aDNA discussed possible leaching through sedimentary strata of the DNA itself, yet it was typically considered not an issue as most of these studies were conducted under stable permafrost or similar conditions (e.g., Hansen et al., 2006; Haile et al., 2007, but see Andersen et al. 2012). Permafrost thawing and re-deposition of material adds a new dimension to this problem of temporal interpretation. The fact that we retrieved mammoth sequences from both cores of two lakes located approximately 5 km apart suggests that this is not an isolated phenomenon but occurs on a regional or even larger scale. Given the wide spread of abrupt permafrost thaw processes in the Arctic plains (West Siberia, Taimyr, Chukotka, Alaska, Canadian Arctic; Kizyakov et al., 2006 and references therein), the phenomena of disturbed stratigraphy of lacustrine sediments can potentially be observed at a pan-Arctic scale. While this indicates that temporal interpretation of sedimentary aeDNA records should be exercised with caution, our study also demonstrates that a careful evaluation of available information on the site and ecosystem in conjunction with the use of independent dating techniques can uncover incongruencies. This is more difficult in older time periods, where artefactual stratigraphies caused by equivalent processes acting for a limited time will not be detected as easily as in our case of long extinct species. The same applies to extant taxa or those which have undergone extinction or extirpation more recently, the presence of which cannot as easily be excluded as in the current example. However, we suggest that the inclusion of robust dating techniques and knowledge of local geophysical processes can provide good arguments to evaluate the reliability of aeDNA records.

## Materials and Methods

### Field sites, DNA isolation and hybridization capture enrichment

In 2019, sediment cores (6 cm diameter; **Table 1**) were retrieved from two lakes (LK-001 and LK-007, respectively, **Table 1**) on the Yamal peninsula, Siberia, using a UWITEC piston corer (UWITEC, Mondsee, Austria). The lakes were located approximately 5 km apart. The cores were transported to the aDNA laboratories of the University of Konstanz, Germany; from lake LK-001, a secondary core was taken which was sliced in the field at 1-cm steps for radiometric dating from 0 to 39.5 cm depth, performed at the Environmental Radioactivity Research Centre, University of Liverpool (Supplement section 9). Additional 14C dating of three specimens of plant remains, extracted at 36.5 cm, 51 cm and 74 cm, was performed (Supplement section 9).

Sedimentary DNA was isolated from 23 samples of core LK-001 and from 16 samples of core LK-007 using commercially available kits with modified protocols (Supplement section 1). The extracts of core LK-001 were subjected to library preparation for capture enrichment. Enrichment probes were designed from mitogenomes of 17 herbivorous mammal species that currently or previously occurred in the Arctic (**Table S2**) and few lichen sequences (**Table S3**). Genomic libraries were produced according to Li et al., 2013 with some modifications (Seeber et al., 2019; Seeber et al. 2023; Supplement section 2). Filtered reads were mapped to mammalian mitogenomes, followed by BLASTn alignment against the complete NCBI nucleotide database and subsequent metagenomic analyses using MEGAN (Huson et al., 2016). Reads assigned to *Mammuthus* were mapped to a complete *M. primigenius* reference mitogenome (NCBI accession NC_007596.2); reads assigned to *Coelodonta antiquitatis* were mapped to the NCBI reference mitogenome NC_012681.1 2. The reads mapped to mammoth from the top three libraries were assigned to haplogroup using mixemt (https://github.com/svohr/mixemt) with a custom-made representative panel of 15 mammoth mitogenomes (**Fig. 2; Table S6**).

#### Conventional PCR, mammal metabarcoding, and ddPCR

Based on the enriched fragments with the highest coverage, PCR primers specific to *M. primigenius* were designed using Geneious Prime 2022.1.1 (Kearse et al., 2012), i.e., mamm801 (5’-CCCATGCAGGAGCTTCTGTAGA-3’) and mamm800r (5’-GACATAGCTGGAGGTTTTATGT-3’) to produce a 121-bp amplicon of the CO1 gene. The specific PCR conditions are described in Supplement section 6. Mammal metabarcoding PCRs were performed on 21 DNA extracts of core LK-001 and 27 samples of core LK-007, with eight independent replicates, each. Each batch of PCRs included one non-template control. Established metabarcoding primers were used (Giguet-Covex et al., 2014), and human blocking primers (Garcés-Pastor et al., 2021) were included. PCR conditions are described in the supplementary material. Sequencing was performed on an Illumina NovaSeq platform, with 2 x 150 reads. The raw data were processed as described in the supplement. We used the sample LK-001_66.5 of core LK-001 which had produced one of the highest *Mammuthus* read counts after enrichment. The specific target amplified by the primer pair were used to design a probe (5’-GGATACTCCTGCAAGGTGAAGTG-3’). With this probe, ddPCR was performed with 24 extracts of core LK-001 and with 30 extracts of core LK-007, with three replicates, each (Supplement section 8).

## Data availability

Raw sequence data of the hybridization capture were made available as an NCBI BioProject (PRJNA933601).

## Acknowledgements

This research was funded through the 2017-2018 Belmont Forum and BiodivERsA joint call for research proposals, under the BiodivScen ERA-Net COFUND program, and with the funding organizations Deutsche Forschungsgemeinschaft (DFG grant EP-98/3-1 to L.S.E.), Agence Nationale de la Recherche (ANR), Research Council of Norway (NFR), the Swedish Research Council for Environment, Agricultural Sciences and Spatial Planning (Formas), Academy of Finland, National Science Foundation (NSF) and the Natural Sciences and Engineering Research Council of Canada (NSERC-CRSNG). Field work was enabled through a grant of the Young Scholar Fund of the University of Konstanz to L.S.E. Y.D. was supported by the RUDN University Strategic Academic Leadership Program. Expedition logistics were supported by the Department of Science and Innovations of Yamalo-Nenets Autonomous Okrug and the Non-profit organization «Russian Center of Arctic Exploration». We thank PD Dr. Elena Marinova for assistance and discussions concerning dating of plant macrofossils and Patrick Bartolin for assistance in the wet lab.

## References

Armbrecht, L.H., Coolen, M.J.L., Lejzerowicz, F., George, S.C., Negandhi, K., Suzuki, Y., et al. (2019) Ancient DNA from marine sediments: Precautions and considerations for seafloor coring, sample handling and data generation. Earth-Science Rev 196: 102887.

Arnold, L.J., Roberts, R.G., MacPhee, R.D.E., Haile, J.S., Brock, F., Möller, P., et al. (2011) Paper II - Dirt, dates and DNA: OSL and radiocarbon chronologies of perennially frozen sediments in Siberia, and their implications for sedimentary ancient DNA studies. Boreas 40: 417–445.

Baulin, V.V., Chekhovskiy, A.L., and Sukhodolskiy, S.E. (1981) Main stages of permafrost development in the North-East of Western part of USSR and Western Siberia. In History of development of permafrost in Eurasia: on the examples of separate regions. Dubikov, G.I.and Baulin, V.V. (eds). Moscow: Nauka, pp. 41–60.

Boere, A.C., Rijpstra, W.I.C., De Lange, G.J., Sinninghe Damsté, J.S., and Coolen, M.J.L. (2011) Preservation potential of ancient plankton DNA in Pleistocene marine sediments. Geobiology 9: 377–393.

Boessenkool, S., Epp, L.S.L.L.S., Haile, J., Bellemain, E., Edwards, M., Coissac, E., et al. (2012) Blocking human contaminant DNA during PCR allows amplification of rare mammal species from sedimentary ancient DNA. Mol Ecol 21: 1806–1815.

Dvornikov, Y.A., Leibman, M.O., Heim, B., Bartsch, A., Haas, A., Khomutov, A.V., et al. (2016) Geodatabase and WebGIS project for long-term permafrost monitoring at the Vaskiny Dachi Research Station, Yamal, Russia. Polarforschung 85: 107–115.

Dvornikov, Y.A., Leibman, M.O., Heim, B., Bartsch, A., Herzschuh, U., Skorospekhova, T.V., et al. (2018) Terrestrial CDOM in lakes of Yamal Peninsula: Connection to lake and lake catchment properties. Remote Sens 10: 167.

Dvornikov, Y.A., Leibman, M.O., Khomutov, A.V., Kizyakov, A.I., Semenov, P.B., Bussmann, I., et al. (2019) Gas-emission craters of the Yamal and Gydan peninsulas: A proposed mechanism for lake genesis and development of permafrost landscapes. Permafr Periglac Process 146–162.

Fordham, D.A., Brown, S.C., Akçakaya, H.R., Brook, B.W., Haythorne, S., Manica, A., Shoemaker, K.T., Austin, J.J., Blonder, B., Pilowsky, J.A., Rahbek, C., Nogues-Bravo, D. (2022) Process-explicit models reveal pathway to extinction for woolly mammoth using pattern-oriented validation. Ecol Letters, 2022;25:125–137.

Fulton, T.L. and Shapiro, B. (2019) Setting Up an Ancient DNA Laboratory. Ancient DNA. Methods in Molecular Biology. B. Shapiro, A. Barlow, P. D. Heintzman et al. New York, Humana Press, New York, NY 1963., pp. 1–13.

Garcés-Pastor, S., Coissac, E., and Brown, A. (2021) High resolution ancient sedimentary DNA shows that alpine plant biodiversity is a result of human land use. 1–29.

Giguet-Covex, C., Pansu, J., Arnaud, F., Rey, P.-J., Griggo, C., Gielly, L., et al. (2014) Long livestock farming history and human landscape shaping revealed by lake sediment DNA. Nat Commun 5: 3211.

Graham, R.W., Belmecheri, S., Choy, K., Culleton, B.J., Davies, L.J., Froese, D., et al. (2016) Timing and causes of mid-Holocene mammoth extinction on St. Paul Island, Alaska. Proc Natl Acad Sci U S A 113: 9310–9314.

Haile, J., Froese, D.G., MacPhee, R.D.E.E., Roberts, R.G., Arnold, L.J., Reyes, A. V., et al. (2009) Ancient DNA reveals late survival of mammoth and horse in interior Alaska. Proc Natl Acad Sci 106: 22352–22357.

Haile, J., Holdaway, R., Oliver, K., Bunce, M., Gilbert, M.T.P., Nielsen, R., et al. (2007) Ancient DNA Chronology within Sediment Deposits: Are Paleobiological Reconstructions Possible and Is DNA Leaching a Factor? Mol Biol Evol 24: 982–989.

Hansen, A.J., Mitchell, D.L., Wiuf, C., Paniker, L., Brand, T.B., Binladen, J., et al. (2006) Crosslinks Rather Than Strand Breaks Determine Access to Ancient DNA Sequences From Frozen Sediments. Genetics 173: 1175–1179.

Huson, D.H., Beier, S., Flade, I., Górska, A., El-Hadidi, M., Mitra, S., et al. (2016) MEGAN Community Edition - Interactive Exploration and Analysis of Large-Scale Microbiome Sequencing Data. PLOS Comput Biol 12: e1004957.

Kachurin, S.P. (1961) Thermokarst on the USSR territory, Melnikova, N.B. (ed) Moscow: Academy of Sciences USSR.

Kahlke, R.-D.D. and Lacombat, F. (2008) The earliest immigration of woolly rhinoceros (Coelodonta tologoijensis, Rhinocerotidae, Mammalia) into Europe and its adaptive evolution in Palaearctic cold stage mammal faunas. Quat Sci Rev 27: 1951–1961.

Kazantsev, V.S., Krivenok, L.A., and Dvornikov, Y.A. (2020) Preliminary data on the methane emission from lake seeps of the Western Siberia permafrost zone. IOP Conf Ser Earth Environ Sci 606: 012022.

Kearse, M., Moir, R., Wilson, A., Stones-Havas, S., Cheung, M., Sturrock, S., et al. (2012) Geneious Basic: an integrated and extendable desktop software platform for the organization and analysis of sequence data. Bioinformatics 28: 1647–9.

Kizyakov, A.I., Leibman, M.O., and Perednya, D.D. (2006) Destructive relief-forming processes at the coasts of the Arctic plains with tabular ground ice. Kriosf Zemli 10: 79–89.

Kokelj, S.V., Lantz, T.C., Kanigan, J., Smith, S.L., and Coutts, R. (2009) Origin and polycyclic behaviour of tundra thaw slumps, Mackenzie Delta region, Northwest Territories, Canada. Permafr Periglac Process 20: 173–184.

Leibman, M.O. and Kizyakov, A.I. (2007) Cryogenic landslides of Yamal and Yugorskiy peninsulas, Moscow: Rosselkhozakademia.

Li, C., Hofreiter, M., Straube, N., Corrigan, S., and Naylor, G.J.P.P. (2013) Capturing protein-coding genes across highly divergent species. Biotechniques 54: 321–326.

Lord, E., Dussex, N., Kierczak, M., Díez-del-Molino, D., Ryder, O.A., Stanton, D.W.G., et al. (2020) Pre-extinction Demographic Stability and Genomic Signatures of Adaptation in the Woolly Rhinoceros. Curr Biol 30: 3871-3879.e7.

Miller, J.H. and Simpson, C. (2022) When did mammoths go extinct? Nature 612: E1–E3.

Murchie, T.J., Kuch, M., Duggan, A.T., Ledger, M.L., Roche, K., Klunk, J., et al. (2021) Optimizing extraction and targeted capture of ancient environmental DNA for reconstructing past environments using the PalaeoChip Arctic-1.0 bait-set. Quat Res 99: 305–328.

Murchie, T.J., Monteath, A.J., Mahony, M.E., Long, G.S., Cocker, S., Sadoway, T., et al. (2021) Collapse of the mammoth-steppe in central Yukon as revealed by ancient environmental DNA. Nat Commun 12: 1–18.

Nogués-Bravo, D., Rodríguez, J., Hortal, J., Batra, P., and Araújo, M.B. (2008) Climate Change, Humans, and the Extinction of the Woolly Mammoth. PLoS Biol 6: e79.

Parducci, L., Bennett, K.D., Ficetola, G.F., Alsos, I.G., Suyama, Y., Wood, J.R., and Pedersen, M.W. (2017) Ancient plant DNA in lake sediments. New Phytol 214: 924–942.

Rey-Iglesia, A., Lister, A.M., Stuart, A.J., Bocherens, H., Szpak, P., Willerslev, E., and Lorenzen, E.D. (2021) Late Pleistocene paleoecology and phylogeography of woolly rhinoceroses. Quat Sci Rev 263:.

Savvichev, A., Rusanov, I., Dvornikov, Y., Kadnikov, V., Kallistova, A., Veslopolova, E., et al. (2021) The water column of the Yamal tundra lakes as a microbial filter preventing methane emission. Biogeosciences 18: 2791–2807.

Seeber, P.A., McEwen, G.K., Löber, U., Förster, D.W., East, M.L., Melzheimer, J., and Greenwood, A.D. (2019) Terrestrial mammal surveillance using hybridization capture of environmental DNA from African waterholes. Mol Ecol Resour 19: 1486–1496.

Seeber, P.A., Palmer, Z., Schmidt, A., Chagas, A., Kitagawa, K., Marinova-Wolff, E., Tafelmaier, Y. and Epp, L.S., 2023. The first European woolly rhinoceros mitogenomes, retrieved from cave hyena coprolites, suggest long-term phylogeographic differentiation. Biology Letters, 19, p.20230343.

Stefaniak, K., Stachowicz-Rybka, R., Borówka, R.K., Hrynowiecka, A., Sobczyk, A., Moskal-del Hoyo, M., et al. (2021) Browsers, grazers or mix-feeders? Study of the diet of extinct Pleistocene Eurasian forest rhinoceros Stephanorhinus kirchbergensis (Jäger, 1839) and woolly rhinoceros Coelodonta antiquitatis (Blumenbach, 1799). Quat Int 605–606: 192–212.

Stuart, A.J. and Lister, A.M. (2012) Extinction chronology of the woolly rhinoceros Coelodonta antiquitatis in the context of late Quaternary megafaunal extinctions in northern Eurasia. Quat Sci Rev 51: 1–17.

Stuart, A.J., Sulerzhitsky, L.D., Orlova, L.A., Kuzmin, Y. V., and Lister, A.M. (2002) The latest woolly mammoths (Mammuthus primigenius Blumenbach) in Europe and Asia: a review of the current evidence. Quat Sci Rev 21: 1559–1569.

Vohr, S.H., Gordon, R., Eizenga, J.M., Erlich, H.A., Calloway, C.D. and Green, R.E., 2017. A phylogenetic approach for haplotype analysis of sequence data from complex mitochondrial mixtures. Forensic Science International: Genetics, 30, 93–105.

Wang, Y., Pedersen, M.W., Alsos, I.G., De Sanctis, B., Racimo, F., Prohaska, A., et al. (2021) Late Quaternary dynamics of Arctic biota from ancient environmental genomics. Nature 600: 86–92.

