## Supplementary material for "Mitochondrial genomes of Pleistocene megafauna retrieved from recent sediment layers of two Siberian lakes": S 1

### 1. Sampling and DNA isolation

The sediment cores were subsampled according to procedures described by Epp et al., (2019), extracting approximately 1­­–2 g sediment from the center of the core at close depth intervals (**Table S1**) . Sampling was performed in a window-less cold room at 4 °C, adhering to standard aDNA precautions.DNA was extracted using two, highly similar kits. Sample depths and respective extraction protocols for both cores are listed below (**Table S1**).

**Table S1.** Sample depths and extraction protocols.

| a) LK-001 | |  | b) LK-007 | |
| --- | --- | --- | --- | --- |
| Sample depth (cm) | Extraction protocol* |  | Sample depth (cm) | Extraction protocol* |
| 1.5 | PowerLyzer |  | 2.0 | PowerLyzer |
| 4.0 |  |  | 4.0 | PowerSoil Pro |
| 7.0 |  |  | 6.0 | PowerLyzer |
| 11.0 |  |  | 8.0 | PowerSoil Pro |
| 12.0 |  |  | 10.0 | PowerLyzer |
| 15,5 |  |  | 12.0 | PowerSoil Pro |
| 18.0 |  |  | 14.0 | PowerSoil Pro |
| 21.5 |  |  | 18.0 | PowerLyzer |
| 23.0 |  |  | 20. | PowerSoil Pro |
| 28.0 |  |  | 24.0 | PowerLyzer |
| 31.5 |  |  | 26.0 | PowerSoil Pro |
| 35.0 |  |  | 30.0 | PowerSoil Pro |
| 40.0 |  |  | 32.0 | PowerSoil Pro |
| 44.0 |  |  | 36.0 | PowerLyzer |
| 46.5 |  |  | 38.0 | PowerSoil Pro |
| 49.0 |  |  | 42.0 | PowerSoil Pro |
| 51.0 |  |  | 44.0 | PowerLyzer |
| 62.0 |  |  | 48.0 | PowerLyzer |
| 65.5 |  |  | 50.0 | PowerLyzer |
| 66.5 |  |  | 54.0 | PowerLyzer |
| 69.5 |  |  | 56.0 | PowerLyzer |
| 73.5 |  |  | 60.0 | PowerLyzer |
| 80.0 |  |  | 62.0 | PowerSoil Pro |
|  |  |  | 64.0 | PowerLyzer |
|  | |  | 66.0 | PowerSoil Pro |
|  |  |  | 68.0 | PowerLyzer |
|  |  |  | 70.0 | PowerSoil Pro |

* Kits from Qiagen (Hilden, Germany)

The PowerLyzer DNA extraction was performed according to a previous study (Alsos et al., 2020), with some modifications:

| *Day 1:* | 1) add 750 μL PowerBead Solution to the PowerBead Tube  2) transfer 0.25–0.35 g manually homogenized sediment to PowerBead Tube  3) FastPrep bead beating: two times Quickprep protocol (20 s at 4.0 m/s); briefly centrifuge to eliminate foam  4) Lysis mix (per sample):  Solution C1 60 µL  Proteinase K (20 mg/mL) 2 µL  1 M DTT 25 µL  Vortex and briefly centrifuge.  7) Add 87 μL lysis mix to each PowerBead Tube; vortex for 5 min; invert the tube, flick and vortex to dissolve pellet, if present.  8) Incubate overnight at 56 °C and 12 rpm |
| --- | --- |
| *Day 2:* | 1) Remove PowerBead Tube from the incubator oven and allow to cool to room temperature  2) Centrifuge at 10,000 x *g* for 1 min  3) Add 250 μL Solution C2 to a 2-mL collection tube  4) Avoiding the pellet, pour the supernatant from step 2 into the collection tube containing Solution C2; vortex for 5 s  5) Incubate at 2–8 °C for 10 min  6) Centrifuge at 10,000 x *g* for 1 min  7) Label a new clean 2 mL collection tube and add 250 μL Solution C3  8) Avoiding the pellet, pour up to 800 μL of supernatant to the collection tube containing Solution C3; vortex for 5 s  9) Incubate at 2–8 °C for 10 min  10) Centrifuge at 10,000 x *g* for 1 min  11) Label a new clean 5 mL collection tube and add 1,400 μL Solution C4  12) Avoiding the pellet, pour 880 μL of supernatant to the collection tube containing Solution C4; vortex for 5 s;  briefly centrifuge  13) Load 650 μL of the Solution C4-supernatant mix on a Spin Column  14) Centrifuge the Spin Column at 10,000 x *g* for 1 min  15) Transfer the Spin Column to a new 2 mL collection tube  16) Repeat steps above until all Solution C4-supernatant mix has been loaded onto the Spin Column  17) Centrifuge the Spin Column at 10,000 x *g* for 1 min  18) Load 500 μL of Solution C5 on the Spin Column  19) Centrifuge the Spin Column at 10,000 x *g* for 1 min  20) Transfer the Spin Column to a new 1.5 mL collection tube  21) Centrifuge the Spin Column at 10,000 x *g* for 1 min  22) Transfer the Spin Column to a new, labelled, and sterile 1.5 mL collection tube  23) Add 65 μL of elution buffer to the center of the filter membrane  24) Incubate at room temperature for 10 min  25) Centrifuge the Spin Column at 10,000 x *g* for 1 min  26) Repeat steps above for a final elution volume of 120–125 μL |

PowerSoil Pro DNA extraction was performed as follows:

| *Day 1:* | 1) Spin the PowerBead Pro Tube briefly; add up to 500 mg soil and 800 μL Solution CD1. Vortex briefly to mix. Add 20 µL Proteinase K (2mg/mL). 2) FastPrep bead beating: two times 20 s at 4.0 m/s; briefly centrifuge to eliminate foam. Incubate at 56 °C overnight. |
| --- | --- |
| *Day 2:* | 3) Centrifuge the PowerBead Pro Tube at 15,000 x *g* for 1 min. 4) Transfer the supernatant to a clean 2 mL microcentrifuge tube. 5) Add 200 μL Solution CD2 and vortex for 5 s. 6) Centrifuge at 15,000 x *g* for 1 min at room temperature. Transfer up to 700 μL supernatant to a clean 2 mL microcentrifuge tube. 7) Add 600 μL Solution CD3 and vortex for 5 s. 8) Load 650 μL of the lysate on an MB Spin Column and centrifuge at 15,000 x *g* for 1 min. 9) Discard the flow-through and repeat step 8 to ensure that all of the lysate has passed through the MB Spin Column. 10) Place the MB Spin Column in a clean 2 mL collection tube. 11) Add 500 μL Solution EA to the MB Spin Column. Centrifuge at 15,000 x *g* for 1 min. 12) Discard the flow-through and place the MB Spin Column back into the same collection tube. 13) Add 500 μL Solution C5 to the MB Spin Column. Centrifuge at 15,000 x *g* for 1 min. 14) Discard the flow-through and place the MB Spin Column in a new 2 mL collection tube 15) Centrifuge at 16,000 x *g* for 2 min. Place the MB Spin Column in a new 1.5 mL elution tube. 16) Add 50–100 μL Solution C6 to the center of the white filter membrane; incubate for 5 min, and centrifuge at 15,000 x *g* for 1 min. |

### 2. Preparation of genomic libraries for hybridization capture

Double-stranded libraries were prepared from genomic DNA according to the protocol below and were visualized by gel electrophoresis. Each batch of libraries (N = 12) was processed with a library blank to control for contamination during library prep. Indexing was performed using individual combinations of P5/P7 index primers.

| **End repair** | | per library |
| --- | --- | --- |
| Mix following components in a sterile low-binding PCR tube | |  |
|  | NEBNext End Repair Buffer (10X) | 5 µL |
|  | NEBNext End Repair Enzyme Mix | 2.5 µL |
|  | genomic DNA | 42.5 µL |
| Incubate in a thermal cycler for 30 mins at 20 ˚C.  Purify using QIAquick/MinElute PCR purification kit. Elution: add 32 µl buffer EB and incubate at 37 ˚C for 5 min before spinning down the DNA at 13,000 rpm for 1 min. | | |
| **Adapter ligation** | |  |
| Mix following components in a sterile low-binding PCR tube | |  |
|  | Quick Ligation Reaction Buffer (10X) | 10 µL |
|  | nuclease-free water | 4 µL |
|  | P5/P7 adapter mix (50 µM stock) | 1 µL |
|  | DNA as purified in step above | 30 µL |
|  | Quick T4 DNA ligase | 4.8 µL |
| *final adapter concentrations for ancient samples should be between 0.25-0.5 µM.  **it is vital to add ligase after mixing DNA with adaptors. | |  |
| Incubate for 15 min at 25 °C; purify using a QIAquick/MinElute PCR purification kit. Elution: 42 µL buffer EB and incubate at 37 ˚C for 5 min before spinning down the DNA at 13,000 rpm for 1 min. | |  |
| **Fill-In Reaction** | |  |
| Add the following reagents into a low-binding PCR tube | |  |
|  | ThermoPol Reaction Buffer | 5 µL |
|  | dNTPs (10 mM) | 2 µL |
|  | Bst DNA polymerase | 3 µL |
|  | DNA as eluted above | 40 µL |
| Incubate:  20 mins at 65 °C  20 mins at 80 °C  No purification is needed after this step. | |  |

**Indexing PCR**

| **Reagent** | **µL** |
| --- | --- |
| H_2_O | 3.45 |
| Platinum Hifi Taq Buffer 10X (Thermo Fisher Scientific) | 2.50 |
| dNTPs (25 mM) | 0.25 |
| bovine serum albumin (New England Biolabs) | 1.00 |
| MgSO_4_ (50 mM) | 1.00 |
| Platinum HiFi (5 U/µL; Thermo Fisher Scientific) | 0.20 |
| total | 8.40 |
| Index P5 (10 µM) | 0.80 |
| Index P7 (10 µM) | 0.80 |
| template DNA | 15.00 |

| °C | t |  |
| --- | --- | --- |
| 94 | 1 min |  |
| 94 | 15 sec | 8 cycles |
| 60 | 20 sec |  |
| 68 | 60 sec |  |
| 68 | 3 min |  |
| 20 | store |  |

Thereafter, libraries were pooled (four libraries/pool) and were purified using a MinElute PCR purification kit (Qiagen; elution: 2 x 20 µL). Concentrations were measured using a Qubit device (broad-range assay; Thermo Fisher Scientific).

### 3. Hybridization capture enrichment and sequencing

To design RNA oligonucleotide baits for hybridization capture, we compiled the mitochondrial genome sequences of 17 selected mammal species (comprising 282,168 nucleotides [nt]; **Table S2**) and ITS-1 and ITS-2 sequences of *Cladonia rangiferina* (8,546 and 4,750 nt, respectively; **Table S3**). The compiled mitochondrial sequences were submitted to Daicel Arbor Biosciences for custom design of 70-bp baits at 3-fold tiling, collapsed at 99% identity and 85% overlap, resulting in 9,339 unique baits (747,120 nt).

Each purified library pool containing four libraries was subjected to one hybridization capture reaction, apart from libraries produced from extraction blanks (N = 15), library blanks (N = 15), and PCR non-template controls (N = 30), all of which were pooled in one reaction due to the minute DNA concentrations (mostly below detection range).

Hybridization capture was performed according to the MYbaits protocol v. 5.00 (Daicel Arbor Biosciences) with the following modifications: a total amount of 150 ng baits per capture reaction was used, and library pools were incubated for hybridization with the baits at 58 °C for 24 h. Post-capture, the enriched libraries were removed from the beads and were amplified similarly to the PCR protocol used for the indexing PCR and with primers IS5/IS6 (Illumina, San Diego, CA, USA) and 14 cycles. PCR products were then purified using the MinElute PCR Purification Kit (Qiagen), and the final library concentration was measured on an Agilent Bioanalyzer. For sequencing, the enriched libraries were pooled at equal concentrations, and two library pools (produced from 57 and 110 libraries, respectively) were sequenced on an Illumina NovaSeq platform using three Novaseq SP 2 x 150 flow cells.

**Table S2**. Mitogenome templates of 17 mammal species for design of RNA baits.

| **Order** | **Species** | **NCBI accession** | **# of baits** |
| --- | --- | --- | --- |
| Artiodactyla | *Bison bison* | NC_012346.1 | 446 |
|  | *Bos primigenius* | NC_020746.1 | 456 |
|  | *Saiga tatarica* | NC_013996.1 | 538 |
|  | *Ovis canadensis* | NC_015889.1 | 522 |
|  | *Ovibos moschatus* | NC_020631.1 | 542 |
|  | *Cervus elaphus* | NC_007704.2 | 523 |
|  | *Rangifer tarandus* | NC_007703.1 | 526 |
|  | *Alces alces* | NC_020677.1 | 520 |
|  | *Camelus ferus* | NC_009629.2 | 583 |
| Perissodactyla | *Equus przewalskii* | NC_024030.1 | 575 |
|  | *Coelodonta antiquitatis* | NC_012681.1 | 571 |
| Lagomorpha | *Lepus arcticus* | NC_044769.1 | 586 |
|  | *Ochotona collaris* | NC_003033.1 | 591 |
| Proboscidea | *Mammuthus primigenius* | NC_007596.2 | 588 |
| Eulipotyphla | *Sorex tundrensis* | NC_025327.1 | 584 |
| Rodentia | *Castor canadensis* | NC_033912.1 | 584 |
|  | *Dicrostonyx torquatus* | NC_034646.1 | 575 |
| 9,310 | | | |

**Table S3**. *Cladonia rangiferina* sequences for RNA bait design (shown are the NCBI accessions).

**ITS-1**

MN756840.1; DQ394367.1; JQ695919.1; MK179592.1; KP031549.1; KP001202.1; AF458306.1; KT792792.1; MK811970.1; KT792788.1; MK508944.1; GU169225.1; KP001197.1; KP001201.1; MK812260.1; MK811708.1; KY119381.1; MK508952.1; KT792789.1; EU266113.1; KY266884.1; KP001192.1; KP001190.1; JQ695918.1; KT792790.1; JQ695920.1; KP001191.1; AF458307.1; KP001200.1; MK508943.1; MK812460.1; MK508937.1; KT792791.1

***resulting in 23 baits***

**ITS-2**

KT792789.1; KP001190.1; AF458307.1; JQ695919.1; MK179592.1; DQ394367.1; KP001194.1; KP001193.1; KY266884.1; KP001199.1; KP001192.1; MK300750.1; MN756487.1; MK508937.1; KP001200.1; MK812460.1; MK508943.1; KP001191.1; KP031549.1; KP001202.1; AF458306.1; MK811970.1; KY119381.1; KP001198.1; MK812260.1; MK811708.1; GU169225.1; JQ695918.1; KP001201.1

***resulting in 6 baits***

### 4. Bioinformatics of capture enrichment data

Adapter sequences were removed and sequences were filtered using leeHom with the default ancient DNA settings (Renaud et al., 2014). Duplicates were removed, and filtered reads were mapped against a database of 73 mammal mitogenomes (**Table S4**) using BWA v. 0.7.17 (Li and Durbin, 2009) and were blastn-aligned against the complete NCBI Genbank nucleotide database (Benson et al., 2009); retrieved March 14 2022) with a maximum e-value of 0.01. All blast output files were processed using MEGAN Community Edition 6.19.8 using a weighted LCA algorithm (Huson et al., 2016). aDNA degradation was examined using mapDamage 2.2.1 (Ginolhac et al., 2011) against a reference genome of *M. primigenius* (NC_007596.2). The respective command lines are shown in **Table S5**. The reads blast-assigned to *M. primigenius* were mapped to the complete *M. primigenius* reference mitogenome to produce consensus sequences using Geneious Prime 2023.0.1 (Kearse et al., 2012).

**Table S4**. Reference mitogenomes used for mapping

| NC_020679.1 | *Antilocapra americana* |  | NC_018783.1 | *Equus ovodovi* |
| --- | --- | --- | --- | --- |
| NC_012346.1 | *Bison bison* |  | HM118851.1 | *Equus hemionus* |
| NC_020746.1 | *Saiga tatarica* |  | MK982180.1 | *Equus asinus* |
| NC_013996.1 | *Bos primigenius* |  | NC_012681.1 | *Coelodonta antiquitatis* |
| NC_015889.1 | *Ovis canadensis* |  | NC_007596.2 | *Mammuthus primigenius* |
| NC_020630.1 | *Oreamnos americanus* |  | FR691686.1 | *Castor fiber* |
| NC_020631.1 | *Ovibos moschatus* |  | NC_033912.1 | *Castor canadensis* |
| NC_027233.1 | *Bison priscus* |  | NC_034313.1 | *Dicrostonyx groenlandicus* |
| NC_009629.2 | *Camelus ferus* |  | NC_034646.1 | *Dicrostonyx torquatus* |
| KR822422.1 | *Camelops cf. hesternus* |  | JN181159.1 | *Peromyscus leucopus* |
| NC_013836.1 | *Cervus elaphus xanthopygus* |  | NC_006853.1 | *Bos taurus* |
| NC_007704.2 | *Cervus elaphus* |  | KM093871.1 | *Capra hircus* |
| NC_013840.1 | *Cervus elaphus yarkandensis* |  | NC_015241.1 | *Microtus fortis fortis* |
| KP405229.1 | *Alces alces cameloides* |  | KP200876.1 | *Vulpes lagopus* |
| NC_020677.1 | *Alces alces* |  | HM236180.1 | *Ovis aries* |
| NC_020729.1 | *Odocoileus hemionus* |  | KT448275.1 | *Canis latrans* |
| NC_015247.1 | *Odocoileus virginianus* |  | JN632610.1 | *Capreolus capreolus* |
| NC_007703.1 | *Rangifer tarandus* |  | KJ681493.1 | *Capreolus pygargus* |
| KY987554.1 | *Platygonus compressus* |  | JN632629.1 | *Dama dama* |
| NC_002008.4 | *Canis lupus familiaris* |  | KM982549.1 | *Lynx lynx* |
| NC_009686.1 | *Canis lupus lupus* |  | KP202265.1 | *Panthera pardus* |
| NC_013445.1 | *Cuon alpinus* |  | NC_026460.1 | *Rhinolophus macrotis* |
| NC_026529.1 | *Vulpes lagopus* |  | Y07726.1 | *Ceratotherium simum* |
| NC_028302.1 | *Panthera leo* |  | NC_005089.1 | *Mus musculus* |
| NC_022842.1 | *Panthera onca* |  | AM711900.1 | *Meles meles* |
| NC_010642.1 | *Panthera tigris* |  | KM091450.1 | *Mustela erminea* |
| NC_014456.1 | *Lynx rufus* |  | NC_005358.1 | *Ochotona princeps* |
| NC_020642.1 | *Martes americana* |  | NC_012095.1 | *Sus scrofa domesticus* |
| NC_020641.1 | *Neovison vison* |  | DQ480489.1 | *Canis lupus familiaris* |
| NC_024942.1 | *Mustela nigripes* |  | NC_020670.1 | *Crocuta crocuta* |
| NC_020664.1 | *Martes pennanti* |  | NC_011116.1 | *Arctodus simus* |
| NC_020639.1 | *Mustela nivalis* |  | NC_027963.1 | *Sorex araneus* |
| NC_009685.1 | *Gulo gulo* |  | NC_025327.1 | *Sorex tundrensis* |
| NC_011112.1 | *Ursus spelaeus* |  | KJ397607.1 | *Lepus arcticus* |
| NC_003426.1 | *Ursus americanus* |  | NC_001640.1 | *Equus caballus* |
| NC_003427.1 | *Ursus arctos* |  | NC_024030.1 | *Equus przewalskii* |
| NC_003428.1 | *Ursus maritimus* |  |  |  |

**Table S5**. Command lines and software used for initial mapping, processing, and taxonomic assignment:

\#**adapter trimming, filtering, and merging**: leeHom (Renaud et al., 2014):

src/leeHom -t 120 –ancientdna –auto -fq1 file_R1.fastq.gz -fq2 file_R2.fastq.gz -fqo leehom_out

\#**mapping**: BWA (Li and Durbin, 2009) against 75 mammal mitogenomes (Table S3):

bwa index reference_mitogenomes.fasta

bwa aln reference_mitogenomes.fasta leehom_out.fq.gz -l 16000 -n 0.01 -O 2 -o 2 -t 8 > bwa_out.sai

bwa samse reference_mitogenomes.fasta bwa_out.sai leehom_out.fq.gz > bwa_out.sam

samtools view -q >= 30 -S -b bwa_out.sam > bwa_out.bam

\#**remove duplicates**: samtools (Li and Durbin, 2009)

samtools collate -o bwa_out_col.bam bwa_out.bam

samtools fixmate -m bwa_out_col.bam bwa_out_fixmate.bam

samtools sort -o bwa_out_pos.bam bwa_out_fixmate.bam

samtools markdup bwa_out_pos.bam bwa_out_mark.bam

samtools fastq bwa_out_mark.bam > bwa_out.fastq

\#**fastq to fasta**

sed -n ‘1~4s/^@/>/p;2~4p’ bwa_out.fastq > bwa_out.fasta

\#**alignment**: blastn (Altschul et al., 1990)

blastn -db ncbi_nt -query bwa_out.fasta -evalue 0.01 -out blastn_out.fasta

\#**aDNA damage**: mapDamage (Ginolhac et al., 2011)

bwa index reference.fasta

bwa aln reference.fasta sample.fasta > sample.sai

bwa samse referecne.fasta sample.sai sample.fasta > sample.sam

samtools view -q 25 -S -b sample.sam > sample.bam

mapDamage -i sample.bam -r reference.fasta

mapDamage -d results_ sample/ -y 0.1 --plot-only

mapDamage -i sample.bam -r reference.fasta –rescale

mapDamage -d results_sample / --forward --stats-only -v -r reference.fasta

### 5. Mammoth mitogenome haplogroup identification

The sequences mapping to mammoth from the libraries with the most mammoth reads were remapped to the mammoth reference (NC_007596) with minimap2 (https://github.com/lh3/minimap2) to generate a bam file. A panel of 15 mitogenome reference sequences (see Table S5) exemplifying the diversity of mammoths was used to identify variants representative of each haplogroup. In cases where there are multiple reference genomes for a haplogroup, the consensus variants unique to the haplogroup were used except for haplogroups B and D&E where haplosubgroups were also identified. mixemt (Vohr et al., 2017, https://github.com/svohr/mixemt) was then used to detect haplogroups present in the core and estimate the mixture proportions of these haplogroups, using only mapped reads with a map quality of >30 and with the requirement that 40% of the unique defining variants of each haplogroup must be observed to say it is present.

**Table S6**. Reference mammoth mitogenomes used for panel

| **Clade** | **Haplogroup** | **Subgroup** | **Accession numbers** | **Proportion of reads assigned to haplogroup** |
| --- | --- | --- | --- | --- |
| II | A |  | EU153451, EU153450 | - |
| III | B | B0 | KX027526, KX027531 | - |
|  |  | B1 | KX027526 | 0.1615 |
|  |  | B2 | KX027531 | - |
| I | C |  | KX027498, KX027565, KX027567, JF912200, KX027499, KX027502 | - |
| I | D&E | D0 | DQ316067, EU153454, EU153447, EU153456, EU153449, EU153455,  EU153446 | - |
|  |  | D1 | DQ316067 | - |
|  |  | D2 | EU153454 | - |
|  |  | D3 | EU153447 | 0.2790 |
|  |  | D4 | EU153456 | - |
|  |  | D5 | EU153449 | 0.0765 |
|  |  | D6 | EU153455 | 0.4829 |
|  |  | D7 | EU153446 | - |
| I | F |  | KX027503, KX027512, NC_015529, KX027511, KX027548, KX027547, KX027556, KX027559, KX027550 | - |
| Krestovka mammoth | K |  | PRJEB42269 (European Nucleotide Archive) | - |

### 6. Conventional PCR

The PCR reaction mix comprised 17.05 µL H_2_O, 0.2 µL Platinum Taq DNA-Polymerase High Fidelity (Thermo Fisher Scientific, Waltham, MA, USA), 2.5 µL 10X reaction buffer, 0.25 µL dNTPs (25 mM), 1 µL bovine serum albumin (20 mg/mL; New England Biolabs, Ipswitch, MA, USA), 1 µL MgSO_4_ (50 mM), 1 µL of each primer (mamm801 and mamm800r; 10 µM), and 1 µL DNA extract. Thermocycling was performed as follows:

| 94 °C | 2 min |  |
| --- | --- | --- |
| 94 °C | 30 s | 55 cycles |
| 54 °C | 30 s |  |
| 68 °C | 20 s |  |
| 68 °C | 1 min |  |

Amplification products were visualized by gel electrophoresis and were purified from excised gel bands using a NucleoSpin Gel and PCR Clean-up kit (Macherey-Nagel, Düren, Germany), followed by Sanger sequencing.

### 7. Mammal metabarcoding

We used the primers MamP007F and MamP007R (Giguet-Covex et al., 2014) and blockers (Garcés-Pastor et al., 2021), with the following conditions:

| **Reagent** | **µL** |
| --- | --- |
| H_2_O | 14.85 |
| Platinum Hifi Taq Buffer 10X (Thermo Fisher Scientific) | 2.5 |
| bovine serum albumin (New England Biolabs) | 0.2 |
| MgSO_4_ (50 mM) | 1.00 |
| dNTPs (25 mM) | 0.25 |
| Platinum HiFi (5 U/µL; Thermo Fisher Scientific) | 0.20 |
| Blocking primer R (1 µM) | 0.5 |
| Blocking primer F (1 µM) | 0.5 |
| total | 20.00 |
| template DNA | 3.00 |

| °C | t |  |
| --- | --- | --- |
| 94 | 5 min |  |
| 94 | 30 sec | 40 cycles |
| 50 | 30 sec |  |
| 68 | 30 sec |  |
| 68 | 10 min |  |
| RT | store |  |

PCR products were purified using a MinElute kit (Qiagen). Sequencing was performed on an Illumina NovaSeq platform, with 2 x 150 reads. The raw data were processed as described below.

### 8. Bioinformatics of mammal metabarcoding data

The obtained datasets by Illumina Sequencing were then processed as follows:

| Load data | obi import --quality-sanger file_R1.fastq reads1  obi import --quality-sanger file_R2.fastq reads2 |
| --- | --- |
| Import tags | obi import --ngsfilter-input taglist.txt ngsfile |
| *Align paired-end reads* | obi alignpairedend -R reads2 reads1 aligned_reads |
| *Grep entries whose mode are alignment* | obi grep -a mode:alignment aligned_reads good_sequences |
| *Assign alignments to individual PCRs* | obi ngsfilter -t ngsfile -u unidentified_sequences good_sequences identified_sequences |
| *Filter out sequences* | obi grep -p "sequence['score']> 50" identified_sequences identified_sequences_filtered  obi grep -p "sequence['score_norm']> 0.9" identified_sequences_filtered identified_sequences_filtered_adj |
| *Dereplicate Sequences* | obi uniq -m sample identified_sequences_filtered_adj dereplicated_sequences_filtered |
| *Keep only useful tags* | obi annotate -k COUNT -k MERGED_sample dereplicated_sequences_filtered cleaned_metadata_sequences |
| *Discard sequences that are shorter than 60bp (based on primer pair)* | obi grep -p "len(sequence)>=60 and sequence['COUNT']>=10" cleaned_metadata_sequences denoised_sequences |
| *Clean the sequences from PCR/sequencing errors* | obi clean -s MERGED_sample -r 0.05 -H denoised_sequences cleaned_sequences |
| *Load database* | cp STD_MAM_1.dat.gz ~/edna_LauraB/master/mammalia/database/ |
| *Import it into DMS* | obi import /data/scc/edna/LauraBa/master/mammalia/database/STD_MAM_1.dat.gz database_mam  obi import --embl EMBL embl_refs |
| *Download the taxonomy* | wget https://ftp.ncbi.nlm.nih.gov/pub/taxonomy/taxdump.tar.gz |
| *Import the taxonomy in the DMS* | obi import --taxdump /data/scc/edna/LauraBa/master/mammalia/taxdump.tar.gz taxonomy/my_tax |
| *Cleaning the database with in silico PCR* | obi ecopcr -e 3 -l 50 -L 150 -F CGAGAAGACCCTATGGAGCT -R CCGAGGTCRCCCCAACC --taxonomy taxonomy/my_tax embl_refs mam_refs |
| *Filter sequences* | obi grep --require-rank=species --require-rank=genus --require-rank=family --taxonomy taxonomy/my_tax mam_refs mam_refs_clean |
| *Dereplicate identical sequences* | obi uniq --taxonomy taxonomy/my_tax mam_refs_clean mam_refs_uniq |
| *Add taxid at the family level* | obi grep --require-rank=family --taxonomy taxonomy/my_tax mam_refs_uniq |
| *Build the reference database* | obi build_ref_db -t 0.97 --taxonomy taxonomy/my_tax mam_refs_uniq_clean mam_db_97 |
| *Assign each sequence to a taxon* | obi ecotag -m 0.97 --taxonomy taxonomy/my_tax -R mam_db_97 cleaned_sequences assigned_sequences |
| *Align the sequences* | obi align -t 0.95 assigned_sequences aligned_assigned_sequences |
| *Export tables for downstream data analysis* | obi grep -A SCIENTIFIC_NAME assigned_sequences assigned_for_metabR |
| *Output two tables required by metabaR* | obi annotate -k MERGED_sample assigned_for_metabR assigned_for_metabR_reads_table  obi export --tab-output --output-na-string 0 assigned_for_metabR_reads_table > mam_reads_01.txt  obi annotate --taxonomy taxonomy/my_tax \  --with-taxon-at-rank superkingdom \  --with-taxon-at-rank kingdom \  --with-taxon-at-rank phylum \  --with-taxon-at-rank subphylum \  --with-taxon-at-rank class \  --with-taxon-at-rank subclass \  --with-taxon-at-rank order \  --with-taxon-at-rank suborder \  --with-taxon-at-rank infraorder \  --with-taxon-at-rank superfamily \  --with-taxon-at-rank family \  --with-taxon-at-rank genus \  --with-taxon-at-rank species \  --with-taxon-at-rank subspecies \  assigned_for_metabR assigned_for_metabR_taxInfo  obi annotate \  -k BEST_IDENTITY -k TAXID -k SCIENTIFIC_NAME -k COUNT -k seq_length \  -k superkingdom_name \  -k kingdom_name \  -k phylum_name \  -k subphylum_name \  -k class_name \  -k subclass_name \  -k order_name \  -k suborder_name \  -k infraorder_name \  -k superfamily_name \  -k family_name \  -k genus_name \  -k species_name \  assigned_for_metabR_taxInfo assigned_for_metabR_motus  obi export --tab-output assigned_for_metabR_motus > mam_motus_01.txt |

Further processing of the data sets was done using RStudio.

| *Editing files for metabar* | reads<- dt_reads %>%  dplyr::select(-c("DEFINITION", "NUC_SEQ"))%>%  as.data.frame() %>%  janitor::row_to_names(row_number= 897, remove_rows_above = FALSE, remove_row= TRUE) %>%  mutate_if(is.integer,as.numeric) |
| --- | --- |
| *assign name to first column* | reads <- cbind(rownames(reads),reads)  rownames(reads) <- NULL  colnames(reads) <- c(names(reads))  colnames(reads)[1] <- "pcr_id" |
| *edit the names of the column* | reads$pcr_id = strsplit(reads$pcr_id,"[.]")  reads$pcr_id = sapply(reads$pcr_id,  function(x) x[length(x)])  rownames(reads)<- reads$pcr_id |
| *Organizing the motus table* | motus<- dplyr::select(dt_motus, 'ID', 'NUC_SEQ', 'COUNT','BEST_IDENTITY', 'TAXID', 'SCIENTIFIC_NAME', 'superkingdom_name', 'species_name', 'class_name', 'order_name', 'family_name', 'genus_name', 'kingdom_name', 'phylum_name', 'subphylum_name', 'subclass_name', 'suborder_name')  names(motus)[names(motus) == 'NUC_SEQ'] <- 'sequence' |

### 9. Droplet digital PCR (ddPCR)

ddPCR was performed using a Bio-Rad QX200 system (Bio-Rad Hercules, CA, USA) using the following conditions, and data were produced and analyzed as per the standard instructions of the manufacturer. Droplets were generated by using 21 µL of the PCR mixture and 70 µL ddPCR Droplet Reader Oil (Bio-Rad) according to the manufacturer's instructions. The final volume for PCR amplification was 40 µL and was carried out in a C1000 Touch Thermo cycler (Bio-Rad). The products were analyzed using a QX200 Droplet Reader (Bio-Rad); the threshold was set manually to 3,000.

| **Reagent** | **µL** |
| --- | --- |
| ddPCR Supermix for probes | 11 |
| H2O (DEPC) | 6.8 |
| 20x Target-Primers/Probe (FAM) | 1.1 |
| 20x Target-Primers/Probe (HEX) | 1.1 |
| total | 20 |
| template DNA | 2.0 |

| °C | t |  |
| --- | --- | --- |
| 95 | 10 min |  |
| 94 | 30 sec | 40 cycles |
| 50 | 30 sec |  |
| 60 | 30 sec |  |
| 98 | 10 min |  |
| 14 | store |  |

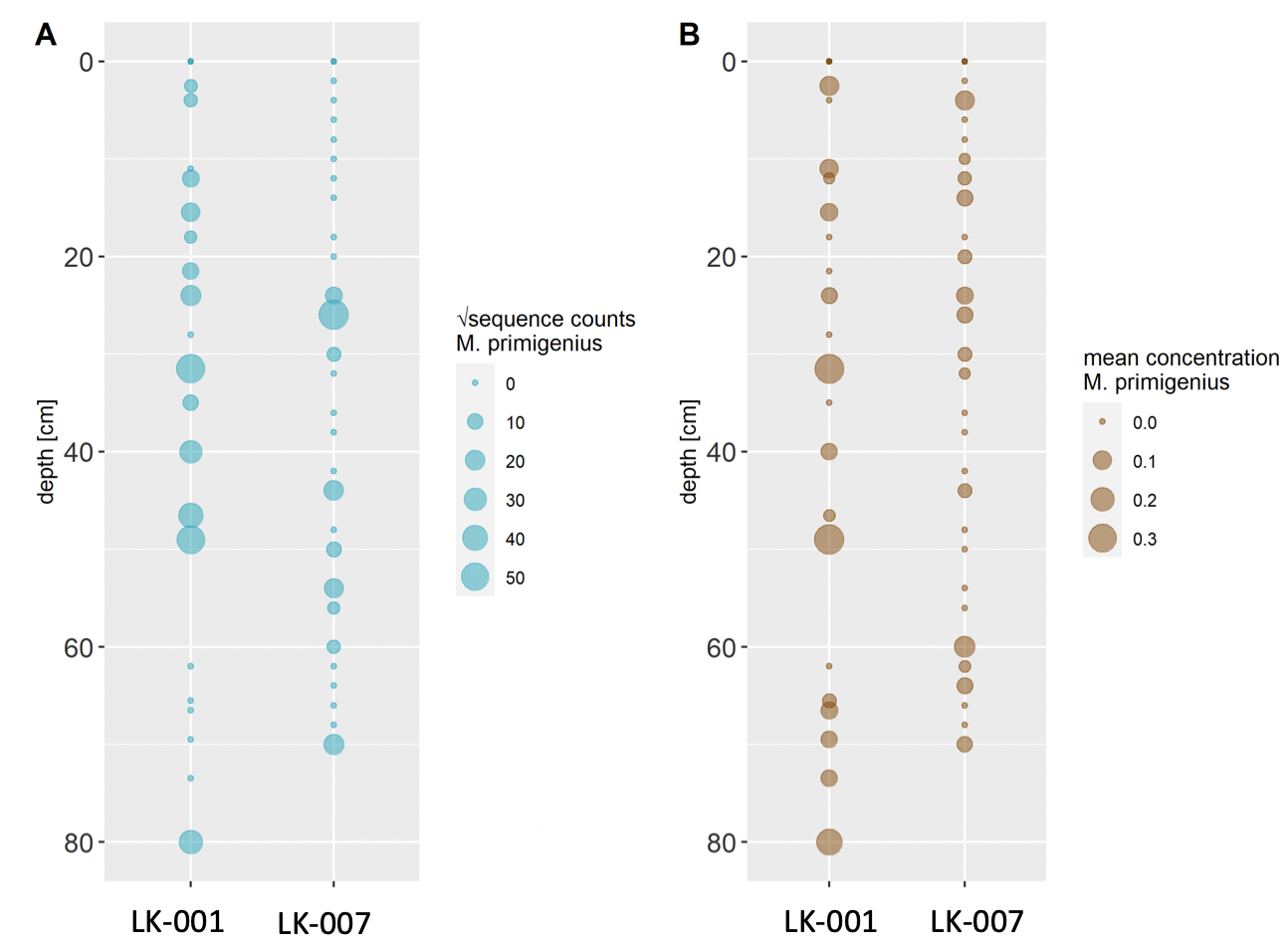

**Fig. S1.** Comparison of transformed metabarcoding read counts (A) and ddPCR concentration estimations (B) of *Mammuthus primigenius* DNA in sediment cores LK-001 and LK-007 from the Yamal peninsula.

### 10. Sediment dating

The dating sections of core LK-001 (**Table S7**) were dried to constant weight at 65 °C and were then analyzed for ^210^Pb, ^226^Ra, and ^137^Cs by direct gamma assay at the Liverpool University Environmental Radioactivity Laboratory using Ortec HPGe GWL series well-type coaxial low background intrinsic germanium detectors. This allowed dating to a depth of 39.5 cm (**Table S7**).

**Table S7**. ^210^Pb chronology of Yamal lake sediment core LK-001.

| Depth | | Chronology | | | Sedimentation Rate | |
| --- | --- | --- | --- | --- | --- | --- |
|  |  | Date | Age |  |  |  |
| cm | g cm^-2^ | AD | y | ± | g cm^-2^ y^-1^ | cm y^-1^ |
| 0.00 | 0.0 | 2019 | 0 | 0 |  |  |
| 0.25 | 0.2 | 2018 | 1 | 1 | 0.34 | 0.37 |
| 1.25 | 1.1 | 2016 | 3 | 2 | 0.34 | 0.37 |
| 2.25 | 1.9 | 2013 | 6 | 2 | 0.34 | 0.37 |
| 3.25 | 2.8 | 2010 | 9 | 2 | 0.34 | 0.37 |
| 4.25 | 3.8 | 2008 | 11 | 2 | 0.34 | 0.37 |
| 5.25 | 4.7 | 2005 | 14 | 2 | 0.34 | 0.37 |
| 6.25 | 5.7 | 2002 | 17 | 2 | 0.34 | 0.37 |
| 7.25 | 6.7 | 1999 | 20 | 3 | 0.36 | 0.37 |
| 8.25 | 7.6 | 1996 | 23 | 3 | 0.39 | 0.41 |
| 10.25 | 9.7 | 1992 | 27 | 3 | 0.49 | 0.47 |
| 12.25 | 11.8 | 1988 | 31 | 4 | 0.53 | 0.49 |
| 14.25 | 14.0 | 1984 | 35 | 4 | 0.55 | 0.51 |
| 14.75 | 14.5 | 1983 | 36 | 4 | 0.55 | 0.51 |
| 15.50 | 15.4 | 1981 | 38 | 4 | 0.55 | 0.51 |
| 16.50 | 16.5 | 1980 | 39 | 4 | 0.55 | 0.51 |
| 17.50 | 17.6 | 1978 | 41 | 4 | 0.55 | 0.51 |
| 18.50 | 18.6 | 1976 | 43 | 4 | 0.55 | 0.51 |
| 20.50 | 20.7 | 1971 | 48 | 5 | 0.55 | 0.51 |
| 22.50 | 22.7 | 1968 | 51 | 6 | 0.55 | 0.51 |
| 23.50 | 23.7 | 1966 | 53 | 6 | 0.55 | 0.51 |
| 24.50 | 24.8 | 1964 | 55 | 6 | 0.55 | 0.51 |
| 25.50 | 25.9 | 1962 | 57 | 7 | 0.44 | 0.42 |
| 26.50 | 27.0 | 1961 | 58 | 7 | 0.37 | 0.36 |
| 27.50 | 28.0 | 1957 | 62 | 8 | 0.30 | 0.29 |
| 28.50 | 29.0 | 1952 | 67 | 9 | 0.28 | 0.22 |
| 29.50 | 30.2 | 1949 | 70 | 10 | 0.23 | 0.19 |
| 31.50 | 32.6 | 1938 | 81 | 10 | 0.23 | 0.19 |
| 33.50 | 34.7 | 1933 | 86 | 10 | 0.23 | 0.19 |
| 35.50 | 36.9 | 1927 | 92 | 11 | 0.23 | 0.19 |
| 37.50 | 39.1 | 1913 | 106 | 13 | 0.23 | 0.19 |
| 39.50 | 41.4 | 1895 | 124 | 17 | 0.23 | 0.19 |

Core LK-001 was additionally dated using radiocarbon. We collected plant remains throughout the core by first using a scalpel to cut 1 cm thick of bulk sediment at desired depths, followed by washing the sediment chunks with distilled water and filtering using 190-µm mesh sieves. We then collected leaves of deciduous trees and seeds, which were placed in glass vials and dried under a fume hood before sending to the Micadas (Alfred Wegener Institute, Bremerhaven, Germany) for radiocarbon dating. Sample preparation and measurement methodologies at the Micadas were performed as described previously (Mollenhauer et al., 2021). We sent three samples that had sufficient plant materials for dating and retained two dates after removing an inaccurate date caused by high inbuilt age from plant remains (**Table S8**).

**Table S8**. Radiocarbon dating results.

| **Sample label** | **F14C** | **± (abs)** | **Age (y)** | **± (y)** | **Weight (µg C)** | **Comment** |
| --- | --- | --- | --- | --- | --- | --- |
| LK-001_36.5cm | 0.9595 | 0.0095 | 332 | 79 | 35 |  |
| LK-001_51cm | 0.3396 | 0.0056 | 8,677 | 132 | 148 | Off; lateral input |
| LK-001_74cm | 0.8248 | 0.0234 | 1,547 | 228 | 13 |  |

### 11. Mammal read numbers

The numbers of reads assigned to all mammals are indicated in **Table S9** (core LK-001 only; hybridization capture experiment).

**Table S9**. Numbers of sequences assigned to mammals (core LK-001).

|  |  | **depth** | | | | | | | | | | | | | | | | | | | | | |
| --- | --- | --- | --- | --- | --- | --- | --- | --- | --- | --- | --- | --- | --- | --- | --- | --- | --- | --- | --- | --- | --- | --- | --- |
|  | **sum** | **1.5** | **4.0** | **7.0** | **11.0** | **12.0** | **15.5** | **18.0** | **21.5** | **23.0** | **28.0** | **31.5** | **35.0** | **40.0** | **46.5** | **49.0** | **51.0** | **62.0** | **65.5** | **66.5** | **69.5** | **73.5** | **80.0** |
| *Mammuthus primigenius* | **19,640** | 191 | 162 | 524 | 290 | 1,205 | 605 | 194 | 666 | 277 | 800 | 3,697 | 305 | 1,188 | 1,374 | 1,923 | 631 | 674 | 519 | 1,110 | 1,083 | 814 | 1,408 |
| *Rangifer tarandus* | **18,055** | 253 | 63 | - | 34 | 328 | 688 | 644 | 1,816 | 545 | 902 | 2,186 | 1,240 | 1,571 | 1,574 | 86 | 284 | 2,714 | 56 | 345 | 991 | 114 | 1,621 |
| *Dicrostonyx torquatus* | **16,870** | 211 | 112 | 34 | 111 | 1,211 | 614 | 271 | 1,035 | 522 | 1,049 | 850 | 1,298 | 314 | 761 | 193 | 318 | 656 | 200 | 2,763 | 1,011 | 1,408 | 1,928 |
| *Lepus* | **6,371** | 148 | 134 | 105 | 23 | 584 | 245 | 40 | 369 | 336 | 630 | 382 | 231 | 404 | 799 | 285 | 141 | 116 | 126 | 352 | 330 | 293 | 298 |
| *Coelodonta antiquitatis* | **2,737** | 33 | - | - | 28 | 119 | 153 | 347 | 55 | - | 97 | 128 | 467 | 298 | 141 | 27 | 103 | 152 | - | 250 | 144 | 4 | 191 |
| *Homo sapiens* | **387** | - | - | - | - | - | - | 38 | 217 | - | - | 23 | - | 25 | - | 69 | - | 15 | - | - | - | - | - |
| *Ovibos moschatus moschatus* | **145** | - | 69 | - | - | - | - | - | - | - | 38 | - | - | - | 38 | - | - | - | - | - | - | - | - |
| *Castor fiber* | **135** | - | - | - | - | 18 | - | - | - | - | 15 | - | - | - | 27 | 51 | - | 6 | 1 | - | - | - | 17 |
| *Bos* | **105** | - | - | - | - | - | - | - | - | - | - | - | - | - | - | 105 | - | - | - | - | - | - | - |
| *Cervinae* | **92** | - | - | - | - | - | - | - | 33 | - | - | - | - | 31 | 28 | - | - | - | - | - | - | - | - |
| *Saiga tatarica* | **91** | - | - | - | - | - | - | - | - | - | 57 | - | 34 | - | - | - | - | - | - | - | - | - | - |
| *Sus scrofa cristatus* | **85** | - | - | - | - | - | - | - | 34 | - | - | - | - | - | - | 24 | - | - | - | - | 27 | - | - |
| *Ochotona* | **74** | - | - | - | - | - | - | - | - | - | - | - | - | - | - | - | - | - | 14 | - | - | - | 60 |
| *Ovis aries musimon* | **70** | - | - | - | - | 70 | - | - | - | - | - | - | - | - | - | - | - | - | - | - | - | - | - |
| *Bison* | **54** | - | - | - | - | 24 | - | 30 | - | - | - | - | - | - | - | - | - | - | - | - | - | - | - |
| *Crocuta crocuta* | **53** | - | - | - | - | - | - | - | - | - | - | - | - | - | - | - | - | - | - | 53 | - | - | - |
| *Chrotopterus auritus* | **43** | - | - | - | - | - | 43 | - | - | - | - | - | - | - | - | - | - | - | - | - | - | - | - |
| *Sorex tundrensis* | **32** | - | - | - | - | - | - | - | - | - | - | - | - | - | - | - | 32 | - | - | - | - | - | - |
| *Equus* | **32** | - | - | - | - | - | - | - | - | - | - | - | - | - | - | - | - | - | 7 | - | - | - | 25 |
| *Peromyscus maniculatus bairdii* | **29** | - | - | - | - | - | - | - | - | - | - | - | - | - | 29 | - | - | - | - | - | - | - | - |
| *Murinae* | **28** | - | - | - | - | - | - | - | - | - | - | - | - | - | - | - | - | - | - | - | - | 28 | - |
| *Capra* | **21** | - | - | - | - | - | - | - | - | - | - | - | - | - | - | - | 21 | - | - | - | - | - | - |
| *Myopus schisticolor* | **18** | - | - | - | - | - | - | - | 18 | - | - | - | - | - | - | - | - | - | - | - | - | - | - |
| *Lemmus trimucronatus* | **16** | 16 | - | - | - | - | - | - | - | - | - | - | - | - | - | - | - | - | - | - | - | - | - |

**References**

Alsos, I.G., Sjögren, P., Brown, A.G., Gielly, L., Merkel, M.K.F., Paus, A., Lammers, Y., Edwards, M.E., Alm, T., Leng, M., Goslar, T., Langdon, C.T., Bakke, J., van der Bilt, W.G.M., 2020. Last Glacial Maximum environmental conditions at Andøya, northern Norway; evidence for a northern ice-edge ecological “hotspot.” Quat. Sci. Rev. 239, 106364. https://doi.org/10.1016/j.quascirev.2020.106364

Altschul, S.F., Gish, W., Miller, W., Myers, E.W., Lipman, D.J., 1990. Basic local alignment search tool. J. Mol. Biol. 215, 403–410. https://doi.org/10.1016/S0022-2836(05)80360-2

Benson, D.A., Karsch-Mizrachi, I., Lipman, D.J., Ostell, J., Sayers, E.W., 2009. GenBank. Nucleic Acids Res. 37, D26–D31. https://doi.org/10.1093/nar/gkn723

Epp, L.S., Zimmermann, H.H., Stoof-Leichsenring, K.R., 2019. Sampling and Extraction of Ancient DNA from Sediments, in: Shapiro, B., Barlow, A., Heintzmann, P., Hofreiter, M., Paijmans, J., Soares, A. (Eds.), Ancient DNA. Methods in Molecular Biology. Humana Press, New York, pp. 31–44. https://doi.org/10.1007/978-1-4939-9176-1_5

Garcés-Pastor, S., Coissac, E., Brown, A., 2021. High resolution ancient sedimentary DNA shows that alpine plant biodiversity is a result of human land use 1–29.

Giguet-Covex, C., Pansu, J., Arnaud, F., Rey, P.-J., Griggo, C., Gielly, L., Domaizon, I., Coissac, E., David, F., Choler, P., Poulenard, J., Taberlet, P., 2014. Long livestock farming history and human landscape shaping revealed by lake sediment DNA. Nat. Commun. 5, 3211. https://doi.org/10.1038/ncomms4211

Ginolhac, A., Rasmussen, M., Gilbert, M.T.P., Willerslev, E., Orlando, L., 2011. mapDamage: testing for damage patterns in ancient DNA sequences. Bioinformatics 27, 2153–2155. https://doi.org/10.1093/bioinformatics/btr347

Huson, D.H., Beier, S., Flade, I., Górska, A., El-Hadidi, M., Mitra, S., Ruscheweyh, H.-J., Tappu, R., 2016. MEGAN Community Edition - Interactive Exploration and Analysis of Large-Scale Microbiome Sequencing Data. PLOS Comput. Biol. 12, e1004957. https://doi.org/10.1371/journal.pcbi.1004957

Kearse, M., Moir, R., Wilson, A., Stones-Havas, S., Cheung, M., Sturrock, S., Buxton, S., Cooper, A., Markowitz, S., Duran, C., Thierer, T., Ashton, B., Meintjes, P., Drummond, A., 2012. Geneious Basic: an integrated and extendable desktop software platform for the organization and analysis of sequence data. Bioinformatics 28, 1647–9. https://doi.org/10.1093/bioinformatics/bts199

Li, H., Durbin, R., 2009. Fast and accurate short read alignment with Burrows-Wheeler transform. Bioinformatics 25, 1754–1760. https://doi.org/10.1093/bioinformatics/btp324

Mollenhauer, G., Grotheer, H., Gentz, T., Bonk, E., Hefter, J., 2021. Standard operation procedures and performance of the MICADAS radiocarbon laboratory at Alfred Wegener Institute (AWI), Germany. Nucl. Instruments Methods Phys. Res. Sect. B Beam Interact. with Mater. Atoms 496, 45–51. https://doi.org/10.1016/j.nimb.2021.03.016

Renaud, G., Stenzel, U., Kelso, J., 2014. leeHom: adaptor trimming and merging for Illumina sequencing reads. Nucleic Acids Res. 42, e141–e141. https://doi.org/10.1093/nar/gku699

Seeber, P.A., Palmer, Z., Schmidt, A., Chagas, A., Kitagawa, K., Marinova-Wolff, E., Tafelmaier, Y. and Epp, L.S., 2023. The first European woolly rhinoceros mitogenomes, retrieved from cave hyena coprolites, suggest long-term phylogeographic differentiation. *Biology Letters*, 19, p.20230343.

Vohr, S.H., Gordon, R., Eizenga, J.M., Erlich, H.A., Calloway, C.D. and Green, R.E., 2017. A phylogenetic approach for haplotype analysis of sequence data from complex mitochondrial mixtures. *Forensic Science International: Genetics*, *30*, 93-105.
